# A novel resistant starch supplement induced significant weight loss in obese/overweight dogs

**DOI:** 10.1101/2025.08.20.671242

**Authors:** M. Davies, D.S. Gardner

**Affiliations:** Provet Limited, Kingsweir, Shaftesbury Road, Gillingham, Dorset SP8 4LJ; School of Veterinary Medicine and Science, University of Nottingham, Sutton Bonington Campus, Leicestershire, LE12 5RD

**Keywords:** Obesity, overweight, canine, dog, resistant-starch, beta-glucan, alpha-lipoic acid, probiotic, weight loss, body condition score

## Abstract

**Objectives:** To evaluate whether a resistant-starch containing supplement administered for 30 days would induce significant weight loss and lower body condition score (BCS) in overweight/obese dogs, compared to placebo.

**Methods:** 21 home-based dogs in Norway completed the study. Study population included 15 breeds of which n=11 were male and n=10 female, ages ranged from 3 – 10 years (Mean 6.7). Weight and BCS were recorded at baseline (Day 0) and after 30 days feeding supplement A (‘active ingredient’). Subsequently, after a 5-day wash-out period with no supplement, all dogs received supplement B (‘placebo’) for a further 30 days, when they were re-weighed and BCS scored. Owners and the veterinarian overseeing the study were blinded to content of supplements.

**Results:** The resistant-starch-containing supplement led to statistically significant weight loss (0.45 ± 0.57 kg, mean ± 1SD; *P* = 0.002) with 17 of 21 dogs (77%) losing weight. A 0.5 unit reduction in BCS was noted (*P* = 0.009). Dogs lost some weight on placebo, but BCS did not change.

**Clinical significance:** This novel supplement containing resistant-starch, beta-glucan, alpha-lipoic acid and probiotic can be used to induce weight loss in overweight/obese dogs.

## INTRODUCTION

Canine obesity is a significant problem worldwide (https://www.worldpetobesity.org/). Dietary fibre is beneficial in managing canine obesity (Chandler 2012). It is postulated that resistant starch may also facilitate weight loss in overweight dogs (Cho et al. 2023; Seo et al. 2022; Beloshapka, Cross, and Swanson 2021). Beta-glucan has also been shown to induce positive metabolic changes in obese dogs (Amaral et al. 2020; Ferreira et al. 2022), while L-carnitine may affect fat metabolism to facilitate weight loss (Sunvold et al. 1998; Blanchard et al. 2004), as has been shown in cats (Center et al. 2012). Alpha-lipoic acid is an effective antioxidant when administered to dogs (Anthony et al. 2021). Free radical accumulation and chronic inflammation can be a feature of overweight and obese dogs. Probiotics have been shown to alter the gut microbiome in the context of obesity, resulting in weight loss (Kang et al. 2024).

The aim of this prospective, masked, cross-over study was to establish whether a unique supplement containing resistant-starch, beta-glucan, alpha-lipoic acid, L-carnitine and a probiotic would cause weight loss in overweight/obese dogs. The null hypothesis was that this resistant-starch containing supplement administered for a 30-day period would not cause significant weight loss in overweight/obese dogs.

## MATERIALS AND METHODS

The study was conducted in accordance with ethical guidelines from the Royal College of Veterinary Surgeons on clinical research in practice (Royal College of Veterinary Surgeons 2024) and was overseen by a practising veterinary surgeon. 27 home-based, client-owned dogs were recruited into the study through a Veterinary Practice based in Norway. 15 different breeds (one crossbreed) were represented from small (e.g. terriers), medium (e.g. sheepdog) to large (e.g. Portuguese water dog) breed dogs. All clients gave consent for the study after reading and signing a participant information sheet. A clinical examination by a veterinarian was conducted before inclusion in the study. The same veterinarian oversaw the dogs’ health throughout and was responsible for weighing and assessing body condition score (Figure 1). Initial body weight ranged from 5.7 to 38.4 kg; initial BCS was 8 (min of 7 to max of 9; 13 of 21 dogs were considered ‘clinically obese’ having a BCS of ≥8). The age range was from 3 – 10 years. The ratio of males: females was 11:9 (55% male).

**Figure 1.**
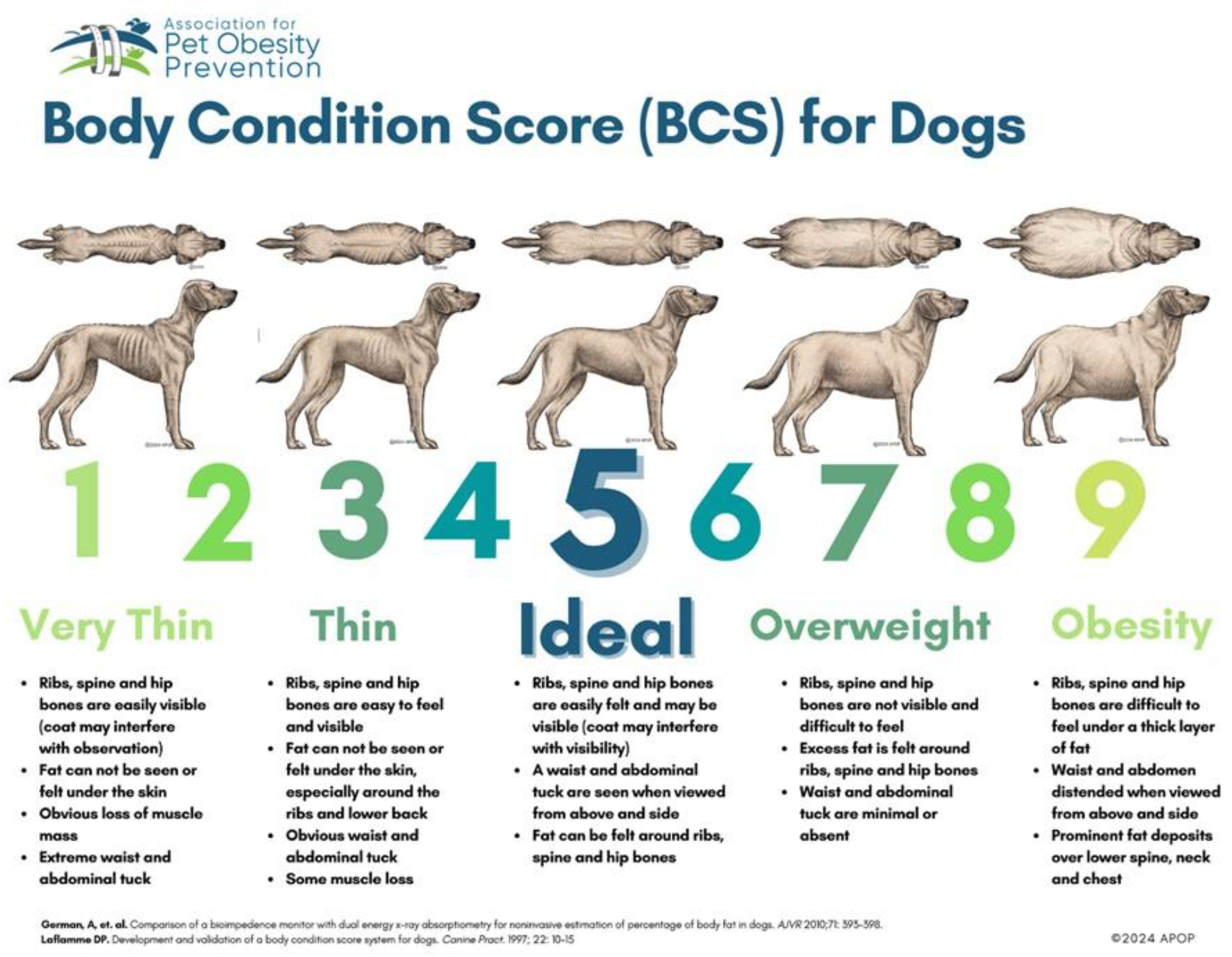
Validated 9-point body condition score chart

Throughout the study all dogs remained in their home environment, were exercised as usual and were fed their usual ration. All dogs were weighed at baseline (Day 0) and after 30-days feeding ‘Supplement A’ (containing novel, active ingredients). After a 5-day wash-out period, all dogs received ‘Supplement B’ (placebo – containing coconut, brewers yeast and a coral palatant) for a further 30 days, at which time they were re-weighed and BCS re-assessed. Owners and the veterinary surgeon overseeing the study were blinded to whether supplement A or B were active ingredient or placebo. The primary outcome was change in body weight (objective) and subjective assessment of body condition score (according to a validated 9-point scale; see Figure 1). At the end of the trial, all owners completed a survey asking eight questions about feeding practices, appetite, activity level and general indices of health and behaviour (see Table 1).

**Table 1.** Survey questions for each owner.

| Number | Question | Possible responses |
| --- | --- | --- |
| 1 | How many times per day did you feed your dog? | Once/twice/three-times/free (~portion size) |
| 2 | How did your dog accept the food? | Normal/rejected/ate slowly/enthusiastic |
| 3 | What was your dog's daily activity level? | None/normal/active ($\geq 1h$)/very active (2h+) |
| 4 | What was the consistency of your dog's stool? | Normal/hard/loose/soft, well-formed |
| 5 | Did you notice any changes in your dog's general appetite? | Less hungry/more hungry/no change |
| 6 | How much of the usual daily food did your dog eat? | All/most/some/none |
| 7 | How easy was it to give the supplement daily? | very easy/adjusted/difficult |
| 8 | General feedback | free text |

Weight data were analysed by paired t-test (baseline – product), BCS by Kolmogorov-Smirnov two-sample nonparamtric test using Genstat v24 (VSNi Ltd, Rothampsted, UK).

## RESULTS

21 of 27 (78%) dogs completed the study. 6 dogs (22%) were withdrawn either because they refused to eat the supplement (n=3) or for medical reasons unrelated to the study (n=1 had spontaneous seizures unrelated to the product, n=1 found to have a tumour and had to be euthanised, n=1 had surgery for an unrelated problem and the owner withdrew consent).

### Effect on bodyweight

Veterinary assessment after 30 days feeding the supplement indicated that n=17 of 21 dogs (81%) lost weight (mean, 0.45 ± 0.57 kg; median, 0.5 kg, mode, 0.3 kg, range from minimum of 0.1 kg to maximum of 1.8 kg (Figure 2; Table 2), representing a percentage loss (with respect to starting weight) of 2.9 % (1.3 – 4.1 %) median (IQR). In one dog there was no change in weight and 3 dogs (14.3%) gained weight (by a mean of 0.2kg). After 30 days on the placebo, relative to last weight recorded then n=19 dogs (90.5%) lost a mean of 0.48 kg (median, 0.2kg). In n=2 dogs there was no change in weight. The percentage weight loss on placebo was 1.4 (0.8 – 5.2%) median (IQR).

**Table 2.** Body Weight Changes.

| DOG | DAY 0<br>(kg) | DAY 30<br>(kg) | DAY 65<br>(kg) | Weight loss<br>to d30 (kg) | Weight loss<br>d30 to d60 (kg) | CHANGE |
| --- | --- | --- | --- | --- | --- | --- |
| 1. Tassen | 9.1 | 8.7 | 8.6 | -0.4 (-4.4%) | -0.1 (-1.2%) | -0.1 kg |
| 2. Tarzan | 5.0 | 5.8 | 5.5 | 0.1 (-1.7%) | -0.3 (-5.4%) | -0.3 kg |
| 3. Nala | 15.3 | 15.1 | 14.2 | -0.2 (-1.3%) | -0.9 (-6.3%) | -0.9 kg |
| 4. Albus | 8.3 | 8.0 | 7.9 | -0.3 (-3.6%) | -0.1 (-1.3%) | -0.1 kg |
| 5. Vita | 10.2 | 9.9 | 9.7 | -0.3 (-2.9%) | -0.2 (-2.1%) | -0.2 kg |
| 6. Bessie | 8.4 | 8.8 | 7.8 | 0.4 (-4.7%) | -1.0 (-12.8%) | -1.0 kg |
| 7. Isa | 9.4 | 9.7 | 9.0 | 0.3 (-3.2%) | -0.7 (-7.7%) | -0.7 kg |
| 8. Stella | 6.1 | 6.0 | 6.0 | -0.1 (-1.6%) | 0.0 (-0.0%) | 0 |
| 9. Aico | 38.4 | 37.0 | 37.5 | -0.9 (-2.3%) | 0.0 (-0.0%) | 0 |
| 10. Humle | 13.2 | 12.9 | 12.8 | -0.3 (-2.3%) | -0.1 (-0.8%) | -0.1 kg |
| 11. Luddi | 20.9 | 21.0 | 20.4 | 0.1 (-0.5%) | -0.6 (-2.9%) | -0.6 kg |
| 12. Happy | 9.1 | 8.8 | 8.6 | -0.3 (-3.3%) | -0.2 (-2.3%) | -0.2 kg |
| 13. Theo | 22.6 | 21.7 | 21.3 | -0.9 (-3.9%) | -0.4 (-1.8%) | -0.4 kg |
| 14. Zappa | 38.0 | 36.3 | 34.4 | -1.8 (-4.9%) | -1.9 (-5.5%) | -1.9 kg |
| 15. Leo | 32.5 | 30.8 | 29.0 | -1.7 (-5.2%) | -1.8 (-6.2%) | -1.8 kg |
| 16. Ludo | 14.4 | 14.4 | 14.2 | 0.0 (-0.0%) | -0.2 (-1.5%) | -0.2 kg |
| 17. Sonic | 19.8 | 19.2 | 19.1 | -0.6 (-3.0%) | -0.1 (-0.5%) | -0.1 kg |
| 18. Stili | 14.8 | 14.0 | 14.1 | -0.8 (-5.4%) | 0.1 (-0.7%) | -0.1 kg |
| 19. Tiny | 13.9 | 13.1 | 13.0 | -0.8 (-5.7%) | -0.1 (-0.7%) | -0.1 kg |
| 20. Presley | 17.1 | 16.4 | 16.2 | -0.7 (-4.1%) | -0.2 (-1.2%) | -0.2 kg |
| 21. Svala | 12.4 | 12.1 | 12.0 | -0.3 (-2.4%) | -0.1 (-0.8%) | -0.1 kg |

**Figure 2.**
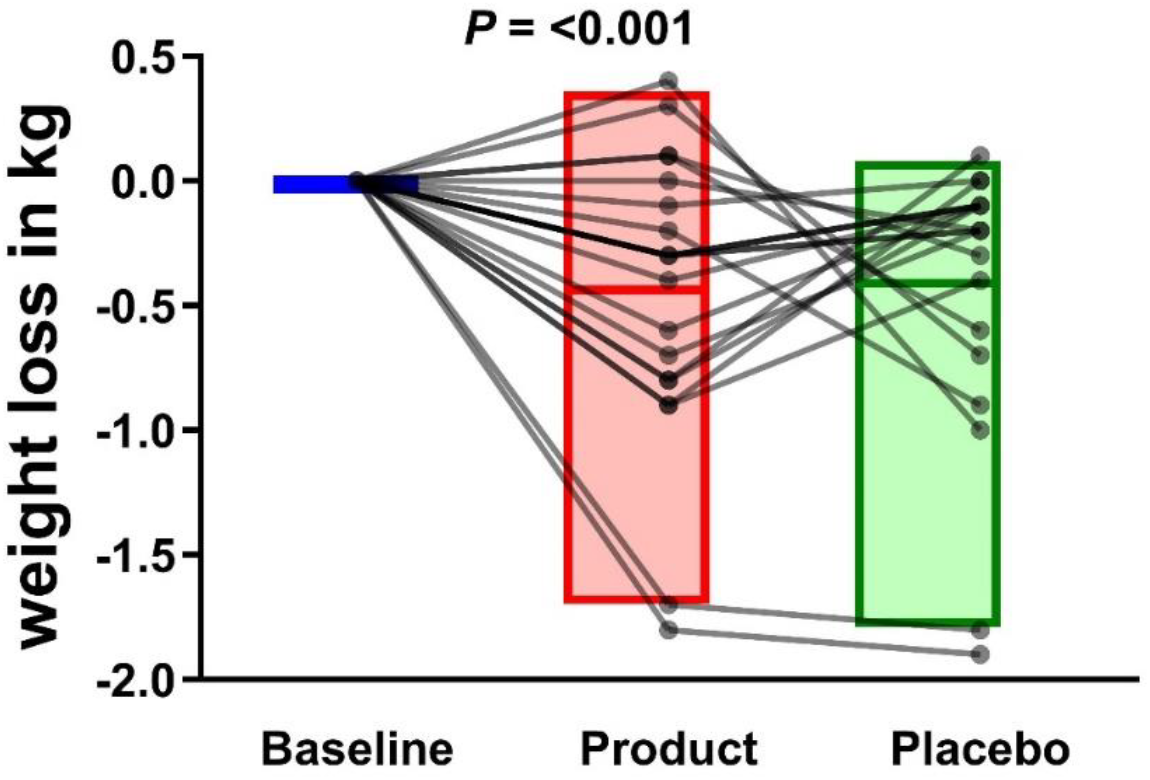
Weight loss on active supplement versus Placebo. **Figure:** Data are change in weight for individual dogs from the beginning of the trial (baseline – adjusted to zero for each dog), after 30-days feeding active supplement (‘Product’), with 5 days wash-out before 30-days feeding placebo (‘Placebo’). Study investigators were blinded to content of each supplement. Consistent weight loss notable on product vs variable response on placebo (see Methods and Results for further detail)

### Effect on BCS

Over 30-days feeding the supplement, body condition score reduced by 0.5 (0 – ^-^1.0) units (median, IQR). 14 of 21 (66%) dogs lost at least 0.5 BCS unit, as assessed by the attending veterinarian. In the remaining n=7 dogs, no change in BCS was recorded, yet these dogs still recorded an average weight loss of 0.41 ± 0.32 kg). When fed placebo, n=8 dogs lost at least 0.5 units BCS, with n=13 of 21 (62%) recording no change in BCS (Table 3).

**Table 3.** Body Condition Score Changes.

| **DOGS** | **DAY 0** | **DAY 30** | **DAY 65** | **Change in BCS**<br>**(0 to 30)** | **Change in BCS**<br>**(30 to 60)** |
| --- | --- | --- | --- | --- | --- |
| 1. Tassen | 8 | 7 | 7 | -1 | 0 |
| 2. Tarzan | 7.5 | 7.5 | 7 | 0 | -0.5 |
| 3. Nala | 7 | 7 | 6 | 0 | -1 |
| 4. Albus | 7.5 | 7 | 7 | -0.5 | 0 |
| 5. Vita | 7.5 | 7 | 7 | -0.5 | 0 |
| 6. Bessie | 8 | 8.5 | 7 | 0.5 | -1.5 |
| 7. Isa | 8 | 8 | 7 | 0 | -1 |
| 8. Stella | 8 | 8 | 8 | 0 | 0 |
| 9. Aico | 9 | 8.5 | 8.5 | -0.5 | 0 |
| 10. Humle | 8.5 | 8 | 8 | -0.5 | 0 |
| 11. Luddi | 9 | 9 | 9 | 0 | 0 |
| 12. Happy | 8 | 8 | 7 | 0 | -1 |
| 13. Theo | 8.5 | 8 | 8 | -0.5 | 0 |
| 14. Zappa | 8 | 7 | 6 | -1 | -1 |
| 15. Leo | 8 | 7 | 6.5 | -1 | -0.5 |
| 16. Ludo | 8 | 8 | 8 | 0 | 0 |
| 17. Sonic | 7.5 | 6.5 | 6 | -1 | -0.5 |
| 18. Stili | 7.5 | 6.5 | 6.5 | -1 | 0 |
| 19. Tiny | 7.5 | 6 | 6 | -1.5 | 0 |
| 20. Presley | 8 | 7 | 7 | -1 | 0 |
| 21. Svala | 7.5 | 7 | 7 | -0.5 | 0 |

### Effect on appetite

All owners of dogs completing the study (n = 21) completed the survey. 18 of 21 owners fed their dogs twice daily. Owners reported that 19 of 21 dogs (90.5%) ate all of their food whilst receiving the active ingredient and that 10 of 21 seemed less hungry (the remaining 11 of 21 reported ‘no change’). This contrasts with only 5 of 21 owners reporting that their dogs seemed less hungry whilst receiving placebo (16 of 21 reported ‘no change’). This effect was statistically significant as analysed by the Chi-square test for association (χ^2^ = 7.22, 1*df*; *P* = 0.007). Owners found the supplement easy to administer, with no change in their dogs general gastrointestinal health (95% reported their dogs having normal faeces whilst on the active ingredient). None of the participant dogs showed any serious adverse events attributed to the products being tested. In addition, some owners specifically noted the following whilst feeding the supplement; ‘seemed happier’ (n=3), ‘shinier coats’ (n=2).

## DISCUSSION

This is the first study to demonstrate the efficacy of a novel resistant-starch containing supplement for helping with weight loss in a varied population of overweight/obese dogs. While the predominant ingredient was resistant-starch, the weight loss effect may be influenced by a combinatorial effect with the other ingredients present (beta-glucan, alpha lipoic acid, probiotics).

Under veterinary supervision, weight loss rates of between 0.5% and 2.0% per week have been reported (Linder and Mueller 2014) with an average, and considered healthy, rate of weight loss being approximately ≤1% per week in pet dogs, considered to be overweight or obese. Faster rates of weight loss can lead to loss of lean body muscle mass. The active-ingredient-containing supplement in this study achieved an average loss of 2.9% initial body weight over 30 days or 0.6 – 0.8% per week, which is close to optimal. The range of percent weight loss in our study was from 0.3 – 1.3% per week, likely influenced by a wide range of initial body sizes recruited to the study. A larger initial sample size would help to establish accurate confidence intervals for weight loss when on the active ingredient.

Nevertheless, some dogs on the placebo supplement also lost body weight. The reason(s) for this are unclear, however, both coconut oil and brewers yeast which were components of the placebo, have been shown to have anti-obesity effects and cause a fall in body fat weight in humans and rats (Chang and Kao 2019; Beegum et al. 2024). It is possible such effects are also noted in canines. The novel resistant-starch containing supplement was palatable, easy to administer and a majority of dogs readily consumed all food. Only 3 of the initial cohort of 27 dogs (11%) were withdrawn for reasons related to palatability or owner compliance. Stools were not altered by the supplement and no serious adverse events were reported attributable to the supplement.

A statistically significant number of owners reported apparently reduced appetite (i.e. their dogs ‘seemed less hungry’) whilst being fed the active ingredient containing supplement. However, all bar one continued to eat all of their main ration. Hence, it is likely that various other food-related ‘begging’ behaviours that were not captured in the survey, were apparent or that the dogs ate their food more slowly. Longer-term, larger clinical trials could tease out precise effects on appetite (documenting amount of food eaten or left), whether initial weight loss stabilises after 30-days and whether dose-response studies might be able to successfully titrate primary outcomes.

## STUDY LIMITATIONS

A larger, complete cross-over design is warranted to tease out any potential carry-over effects of the supplement. A wash-out period of 5-days may not be sufficient to mitigate potential metabolic effects, and weighing of dogs before-after both trials (with double-blinding) is also warranted to control for any possible weight change during the re-alimented washout period. Since coconut and brewers yeast are reported to have anti-obesity effects in other species, it may have been prudent to have used an alternate placebo. Intermediate scoring of BCS (e.g. 8.5), whilst used often in practice by experienced assessors, are not validated on the 9-point scale, and should be avoided. Finally, the majority of dogs (>70%) in the UK are neutered (PDSA 2024), whereas routine neutering is illegal in Norway. Neutering is known to affect body weight gain and predispose to obesity in dogs. Thus, the study may not be representative of the UK dog population, and further work is warranted to validate effects in neutered animals.

## CONCLUSIONS

This is the first study to confirm the efficacy of a novel resistant-starch containing nutritional supplement for reducing body weight and body condition score in overweight/obese dogs, compared to placebo. The supplement was palatable, easy to administer and caused no serious adverse events

## ACKNOWLEDGEMENTS

Project members at Passion4food: Anette Berge, Zahra Salimi, Ole Maribu, Milka Tesla. Veterinarian Ole Maribu.

## FUNDING

Study funded by OMNI Pet food Ltd

## Notes

### Competing Interest Statement

MD is an independent veterinary clinical nutrition consultant that has received fees from pet food companies for consultancy work. All these were unrelated to this manuscript. In the past, MD has received consultancy fees from OMNI Pet Food Ltd for work unrelated to this manuscript.
DSG reports no conflicts of interest.
OMNI Pet Food Ltd funded the study at a contract-research organisation, but had no influence over the reporting of results described herein.

### Summary of Updates

no revisions required, just want to submit on to a journal

